# Area postrema neurons mediate interleukin-6 function in cancer-associated cachexia

**DOI:** 10.1101/2023.01.12.523716

**Authors:** Qingtao Sun, Daniëlle van de Lisdonk, Miriam Ferrer, Bruno Gegenhuber, Melody Wu, Jessica Tollkuhn, Tobias Janowitz, Bo Li

**Affiliations:** Cold Spring Harbor Laboratory, Cold Spring Harbor, NY 11724, USA; Center for Neuroscience, University of Amsterdam, Amsterdam, the Netherlands

**Author notes:** **Correspondence and requests for materials** should be addressed to B.L. These authors contributed equally (Qingtao Sun, Daniëlle van de Lisdonk, Miriam Ferrer).

## Abstract

Interleukin-6 (IL-6) has been long considered a key player in cancer-associated cachexia^1-15^. It is believed that sustained elevation of IL-6 production during cancer progression causes brain dysfunctions, which ultimately result in cachexia^16-20^. However, how peripheral IL-6 influences the brain remains poorly understood. Here we show that neurons in the area postrema (AP), a circumventricular structure in the hindbrain, mediate the function of IL-6 in cancer-associated cachexia in mice. We found that circulating IL-6 can rapidly enter the AP and activate AP neurons. Peripheral tumor, known to increase circulating IL-6^1-5,15,18,21-23^, leads to elevated IL-6 and neuronal hyperactivity in the AP, and causes potentiated excitatory synaptic transmission onto AP neurons. Remarkably, neutralization of IL-6 in the brain of tumor-bearing mice with an IL-6 antibody prevents cachexia, reduces the hyperactivity in an AP network, and markedly prolongs lifespan. Furthermore, suppression of *Il6ra*, the gene encoding IL-6 receptor, specifically in AP neurons with CRISPR/dCas9 interference achieves similar effects. Silencing of Gfral-expressing AP neurons also ameliorates the cancer-associated cachectic phenotypes and AP network hyperactivity. Our study identifies a central mechanism underlying the function of peripheral IL-6, which may serve as a target for treating cancer-associated cachexia.

---

Cancer-associated cachexia is a devastating metabolic wasting syndrome characterized by anorexia, fatigue, and dramatic involuntary bodyweight loss^18,19,24,25^. It affects 50-80% of cancer patients, lowering the quality of life, reducing tolerance to anticancer therapies, and drastically accelerating death^19,20^. The brain is known to have an important role in the pathogenesis of cancer-associated cachexia^16-18^. In particular, recent studies implicate the hypothalamus, parabrachial nucleus, area postrema and other hindbrain structures in the development of cachectic phenotypes in animal models of cancer, such as anorexia, weight loss, and accelerated catabolic processes^26-32^. However, how the brain senses and reacts to peripheral cancers, thereby contributing to the development of cachectic phenotypes, is not well understood.

Possible mediators of cancer-associated cachexia that may act as messengers to engage the brain during cancer progression include tumor-derived factors, metabolites from organs indirectly affected by tumor, and immune or inflammatory factors altered by tumor^16-18,24,33,34^. One such messenger is the pleiotropic cytokine IL-6^18-20,23,24,35,36^. Indeed, elevated levels of circulating IL-6 are associated with cancer progression and cachexia in patients and animal models^1-5,15^. Systemic administration of antibodies against IL-6 or IL-6 receptor shows anticachectic effects in human case reports^6,7,37,38^. Consistently, cancer-associated cachexia in mouse models can be ameliorated by peripheral administration of antibodies against IL-6^8-13^ or IL-6 receptor^14^, or by deletion of the *Il6* gene^9,10^. These findings strongly indicate that IL-6 is a key mediator of cancer-associated cachexia.

Most studies and therapeutic explorations on IL-6 in cancer-associated cachexia has focused on its functions in peripheral organs, including the skeletal muscle, liver, and gut^23^. Although previous studies suggest that IL-6 may also influence brain functions – such as the regulation of food intake^39-41^, fever^42^ and the hypothalamic-pituitary-adrenal (HPA) axis^43^ – how peripheral IL-6 is involved in these functions is unclear. In principle, IL-6 can activate its receptors on the terminals of peripheral nerves, which then transmit the signals to the brain^44^. Alternatively, circulating IL-6 may cross the blood-brain barrier (BBB) or reach circumventricular organs that lack or have a weak BBB, thereby acting within the brain^43,45,46^.

## The area postrema senses circulating IL-6

To determine if circulating IL-6 can enter the brain, we administered biotinylated IL-6 to the venous sinus of mice through retro-orbital injection (Methods). Three hours later, we prepared brain sections from these mice in which the presence of the exogenous IL-6 was examined on the basis of avidin-biotin interactions. In the entire brain, we detected the peripherally administered IL-6 only in the area postrema (AP) (Fig. 1a-d; Extended Data Fig. 1), which is a circumventricular organ located outside of the BBB and has been implicated in nausea and vomiting response to emetic agents entering the circulation^47-50^. Furthermore, immunohistochemistry in the AP revealed that the IL-6 administration markedly increased the expression of Fos (Fig. 1e, f), an immediate early gene product linked to recent neuronal activation^51,52^.

**Figure 1.**
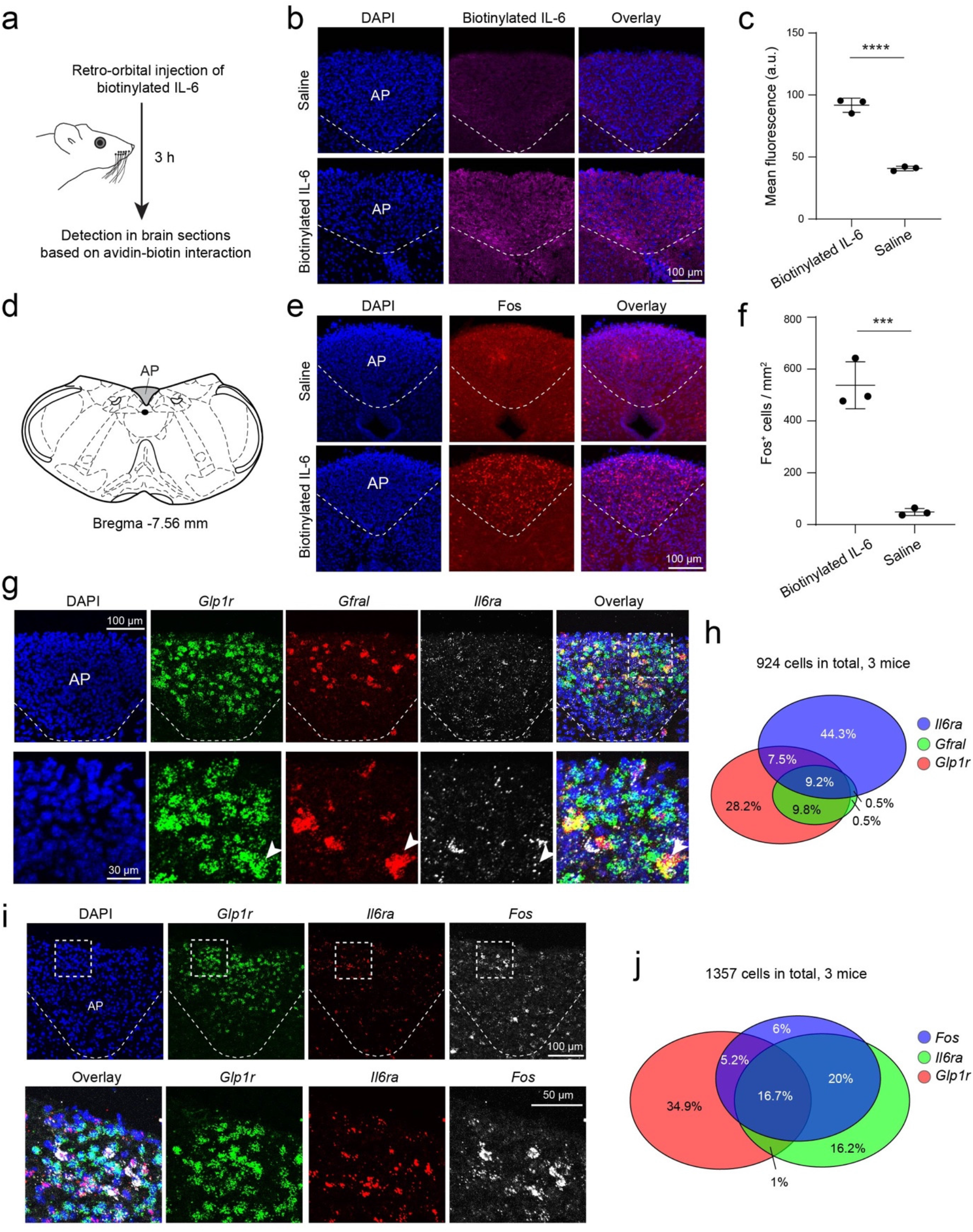
Circulating IL-6 can reach the area postrema (AP) and activate AP neurons. **a**. A schematic of the approach. **b**. Confocal images showing the binding of the exogenous biotinylated IL-6 to cells in the AP. **c**. Quantification of the fluorescence signals from fluorescence-conjugated avidin in the AP, which recognizes the biotinylated exogenous IL-6 (n = 3 mice for each group, t = 24.71, ****P = 0.000016, unpaired t test). **d**. A diagram showing the position of the AP in a coronal brain section. **e**. Confocal images showing Fos expression in the AP. **f**. Quantification of Fos-expressing (Fos^+^) cells in the AP (n = 3 mice for each group, t = 9.246, ***P = 0.0008, unpaired t test). **g**. Confocal images showing the expression of different genes in AP cells, which was detected with single molecule fluorescent *in situ* hybridization (smFISH). At the bottom are higher magnification images of the boxed area in the overlay image on the top. Arrowheads indicate a neuron that expresses all three genes. **h**. A Venn diagram showing the relationships among cells expressing *Il6ra, Gfral*, and *Glp1r* in the AP. **i**. Characterization of the types of *Fos*^*+*^ cells in the AP by smFISH. At the bottom are higher magnification images of the boxed areas in images on the top. **j**. A Venn diagram showing the relationships among cells expressing *Fos, Il6ra*, and *Glp1r* in the AP.

Single molecule fluorescent *in situ* hybridization (smFISH) revealed that the *Il6ra*-expressing (*Il6ra*^+^) cells in the AP partially overlapped with *glucagon-like peptide 1 receptor*-expressing (*Glp1r*^+^) neurons (Fig. 1g-j), the major excitatory neuronal type in the AP^50^. About 17-18% of all the detected AP cells (which likely included glia cells) expressed both *Il6ra* and *Glp1r* (Fig. 1h & j). These *Il6ra*^+^ cells also partially overlapped with *Gfral*-expressing (*Gfral*^+^) neurons (Fig. 1g, h), a subpopulation of *Glp1r*^+^ neurons in the AP that have been implicated in nausea and cancer-associated cachexia^28,29,50^. Interestingly, intravenous IL-6 administration induced *Fos* expression mainly in the *Il6ra*^+^ cells, and these cells partially overlapped with the *Glp1r*^+^ neurons (Fig. 1i, j). These results demonstrate that increased IL-6 in circulation is readily “sensed” by the AP, where it leads to neuronal activation within hours.

### Tumor causes AP hyperactivity

To investigate whether cancer, which is known to increase circulating IL-6^1-5,15^, affects AP neurons, we used mice inoculated with the C26 adenocarcinoma (Fig. 2a, Methods). Mice in this model show persistent increase in blood IL-6 levels, followed by robust cachectic phenotypes, including anorexia and dramatic bodyweight loss^15,18,21-23^. We first measured IL-6 levels in the AP at different timepoints in this model. Notably, IL-6 was increased in the AP on day 7 following tumor inoculation, as well as after the animals had developed cachexia (Fig. 2b).

**Figure 2.**
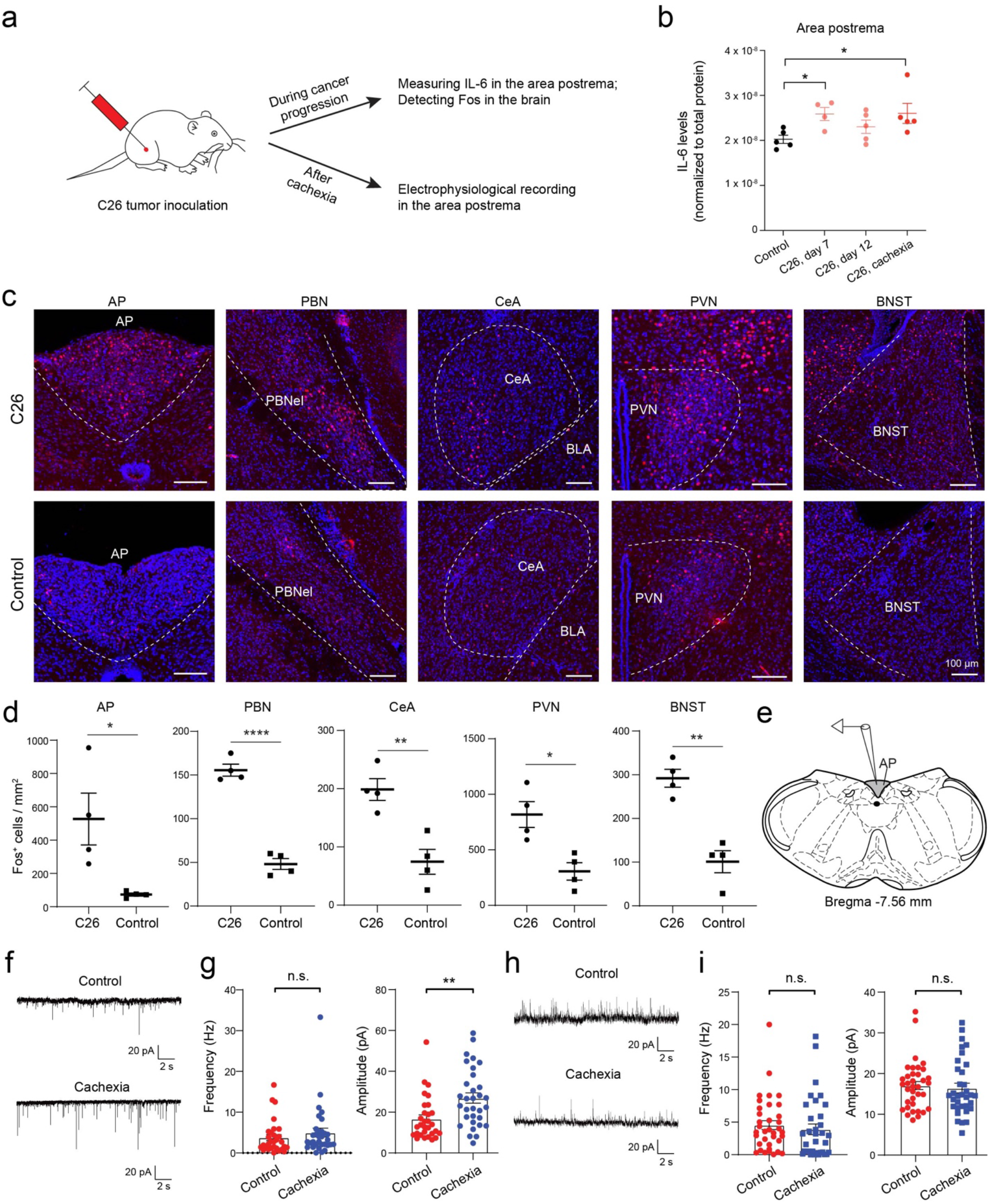
C26 cancer causes increased IL-6 in the AP and AP neuron hyperactivity. **a**. A schematic of the experimental procedure. **b**. IL-6 levels in the area postrema (AP) during cancer progression. IL-6 levels were normalized to total protein levels (n = 4-5 mice in each group, F = 2.9, P = 0.0445, *P < 0.05; One-way ANOVA followed by Tukey’s post-hoc test). **c**. Confocal images showing Fos expression in different brain areas in tumor-bearing (top) or control (bottom) mice. **d**. Quantification of Fos^+^ cells in different brain areas (n = 4 mice in each group; AP, t = 2.91, *P = 0.0268, PBN, t = 11.65, ****P = 2.41 × 10^−5^, CeA, t = 4.37, **P = 0.0047, PVN, t = 3.65, *P = 0.011, BNST, t = 5.86, **P = 0.0011; unpaired t test). **e**. A diagram showing the AP in a coronal brain section for electrophysiological recording. **f**. Representative miniature EPSC traces from AP neurons in control (top) and cachectic (bottom) mice. **g**. Quantification of miniature EPSC frequency (left) and amplitude (right) (control, n = 30 cells / 6 mice, cachexia, n = 32 cells / 7 mice; frequency, n.s. (nonsignificant), P = 0.2513, amplitude, **P = 0.0017, Mann-Whitney test). **h**. Representative spontaneous IPSC traces from AP neurons in control (top) and cachectic (bottom) mice. **i**. Quantification of spontaneous IPSC frequency (left) and amplitude (right) (control, n = 36 cells / 7 mice, cachexia, n = 34 cells / 6 mice; frequency, n.s., P = 0.1637, amplitude, n.s., P = 0.4580, Mann-Whitney test).

We then examined Fos expression in the brain in this model before the onset of cachexia (11 days after tumor inoculation). The number of Fos^+^ cells in the AP was markedly increased in the tumor-bearing mice compared with control mice (Fig. 2c, d). Interestingly, the parabrachial nucleus (PBN), the paraventricular nucleus of the hypothalamus (PVN), the bed nucleus of the stria terminalis (BNST), and the central amygdala (CeA) – which are structures previously implicated in cancer cachexia^17,27-29^ – also showed tumor-induced increase in Fos^+^ cells. Given these structures are interconnected^53-56^ and receive monosynaptic or di-synaptic inputs from the AP^50,57^, these results suggest that cancer progression leads to increased activities in a network of brain areas encompassing the AP.

Next, we tested whether cachexia is associated with lasting functional changes in AP neurons. We prepared acute brain slices from healthy control mice as well as the tumor-bearing mice already showing cachectic phenotypes, and recorded synaptic responses from AP neurons (Fig. 2e). We found that the amplitude of miniature excitatory postsynaptic currents (EPSCs) was markedly increased, while the frequency of the EPSCs was unchanged in cachectic mice relative to control mice (Fig. 2f, g). In contrast, there was no difference in the inhibitory postsynaptic currents (IPSCs) between cachectic mice and control mice (Fig. 2h, i). These results demonstrate that cancer-associated cachexia is accompanied by a potentiation of excitatory synaptic transmission onto AP neurons, which may lead to hyperactivity in these neurons.

### Neutralizing IL-6 in the brain prevents cancer-associated cachexia and AP hyperactivity

We reasoned that increased circulating IL-6 during cancer progression readily enters the AP and results in AP neuron activation – like the intravenously administered exogenous IL-6 (Fig. 1) – leading to hyperactivity in these neurons, which contributes to the development of cachexia.

As a first step to test this hypothesis, we neutralized IL-6 in the brain by intracerebroventricular (i.c.v.) infusion of an antibody against IL-6, which was achieved using an implanted miniature pump (Methods). Continuous infusion of the IL-6 antibody, or an isotype control antibody, was initiated at 10 or 12 days after tumor inoculation (Fig. 3a, b; Extended Data Fig. 2a), a stage when AP neurons show elevated activity (Fig. 2c-d) but cachexia has not yet started. Remarkably, compared with the control antibody, the IL-6 antibody prevented the cachectic phenotypes in almost all the mice, prolonging lifespans (Fig. 3c), reducing bodyweight and tissue loss (Fig. 3d, e; Extended Data Fig. 2b, c), increasing food and water intake (Fig. 3f), and increasing blood glucose levels (Extended Data Fig. 2d). Moreover, IL-6 antibody infusion reduced Fos expression in the AP, PBN, PVN, BNST, and, to a lesser extent, CeA (Fig. 3g, h). As expected, i.c.v. infusion of IL-6 antibodies did not stop tumor from growing (Extended Data Fig. 2e, f).

**Figure 3.**
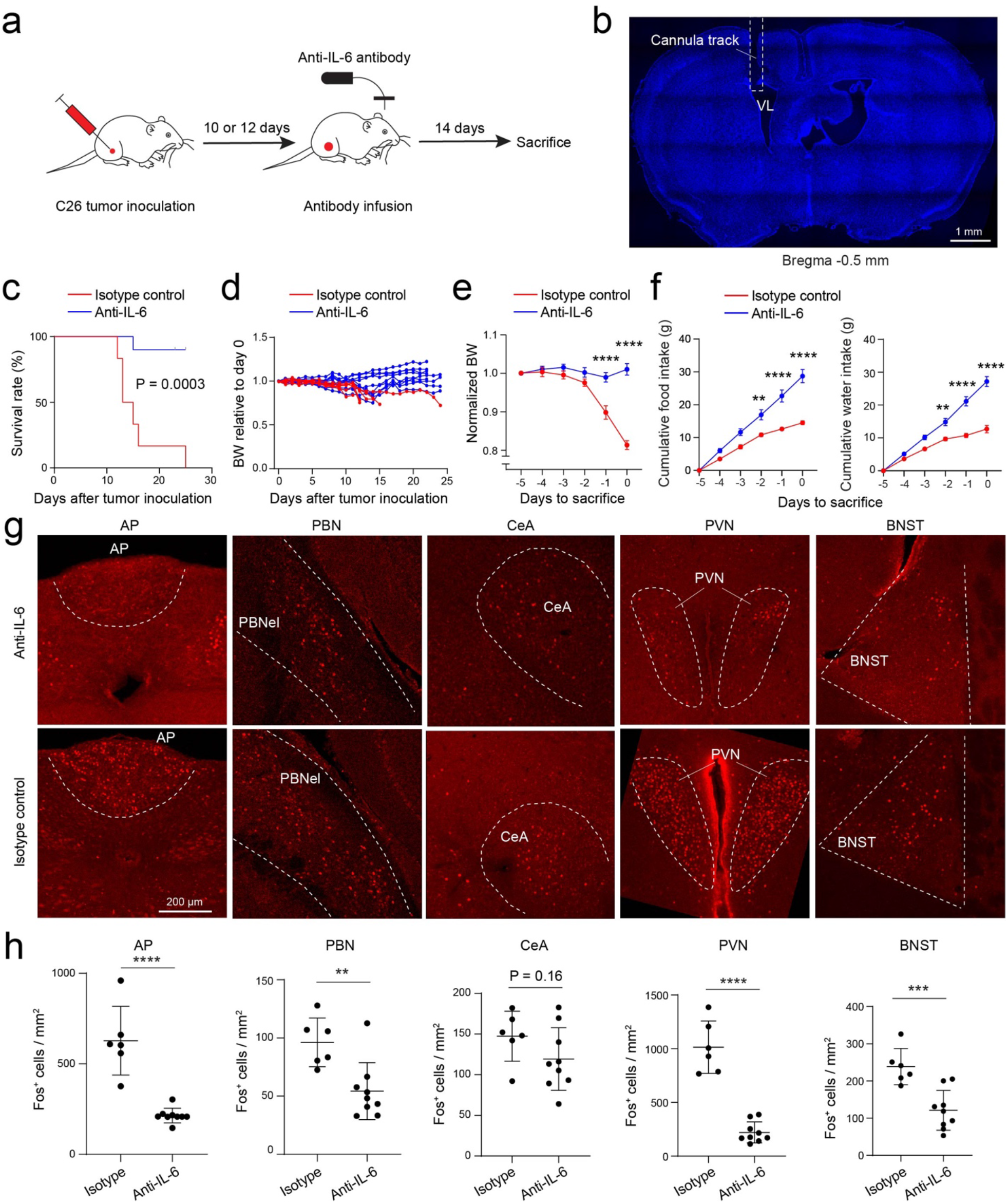
Intracerebroventricular (i.c.v.) infusion of IL-6 antibody prevents cachexia in the C26 cancer model. **a**. A schematic of the experimental procedure. **b**. A confocal image of a coronal brain section from a representative mouse, showing the location of the infusion cannula above the lateral ventricle (VL). **c**. Survival curves of the mice after tumor inoculation (IL-6 antibody group, n = 10, isotype control antibody group, n = 6; P = 0.0003, Mantel-Cox test). **d**. Bodyweight of individual mice relative to their bodyweight on the day of tumor inoculation. **e**. Average bodyweight normalized to that on day -5 (IL-6 antibody, n = 10, isotype control antibody, n = 6; F(1,84) = 75.13, P = 2.77 × 10^−13^; day -1, ****P = 1.2 × 10^−6^, day 0, ****P < 1 × 10^−15^; two-way repeated-measures (RM) ANOVA followed by Sidak’s post hoc test). **f**. Cumulative food (left) and water (right) intake of the mice in the 5 days before sacrifice (IL-6 antibody, n = 10, isotype control antibody, n = 6 mice; food: F(1,84) = 70.87, P = 8.78 × 10^−13^, day -2, **P = 0.007, day -1, ****P = 2 × 10^−6^, day 0, ****P = 8 × 10^−11^; water: F(1,84) = 106.6, P < 1 × 10^−15^, day -2, **P = 0.002, day -1, ****P = 3.6 × 10^−10^, day 0, ****P < 1 × 10^−15^; two-way RM ANOVA with Sidak’s post hoc test). **g**. Confocal images showing Fos expression in different brain areas in the mice infused with the IL-6 antibody (top) and the control antibody (bottom). **h**. Quantification of Fos^+^ cells in different brain areas (IL-6 antibody, n = 9 mice, isotype control antibody, n = 6 mice; AP, t = 6.44, ****P = 2.2 × 10^−5^, PBN, t = 3.43, **P = 0.0045, CeA, t = 1.49, P = 0.16, PVN, t = 8.89, ****P = 6.95 × 10^−7^, BNST, t = 4.309, *** P = 0.0008; unpaired t test).

Thus, reducing IL-6 levels in the brain during cancer progression effectively prevents cachexia, and also dampens the cancer-associated hyperactivity in the AP network.

### Suppression of *Il6ra* expression in AP neurons ameliorates cancer-associated cachexia

To investigate whether AP neurons mediate the functions of IL-6 in the development of cancer-associated cachexia, we sought to suppress *Il6ra*, the gene encoding IL-6Rα, in these neurons using a recently developed CRISPR/dCas9 interference system. This system consists of dCas9-KRAB-MeCP2 [a fusion protein including the nuclease-dead Cas9 (dCas9), a Krüppel-associated box (KRAB) repressor domain, and the methyl-CpG binding protein 2 (MeCP2)] for transcriptional repression, and a CRISPR sgRNA (single guide RNA) for targeting the promoter region of genes of interest^58,59^. We designed and identified two sgRNAs, *Il6ra*-sgRNA-4 and -6, that resulted in suppression of *Il6ra* transcription when co-expressed with dCas9-KRAB-MeCP2 in *in vitro* screening (Extended Data Fig. 3).

We then injected the AP with a lentivirus expressing dCas9-KRAB-MeCP2 in neurons (lenti-SYN-FLAG-dCas9-KRAB-MeCP2), together with a lentivirus expressing *Il6ra*-sgRNA-4 or -6 (lenti-U6-*Il6ra* sgRNA-4/EF1α-mCherry or lenti-U6-*Il6ra* sgRNA-6/EF1α-mCherry, respectively), or a sgRNA targeting the bacterial gene *lacZ* as a control (lenti-U6-*lacZ* sgRNA/EF1α-mCherry)^58^ (Fig. 4a, b). Two weeks later, these mice were inoculated with the C26 tumor.

**Figure 4.**
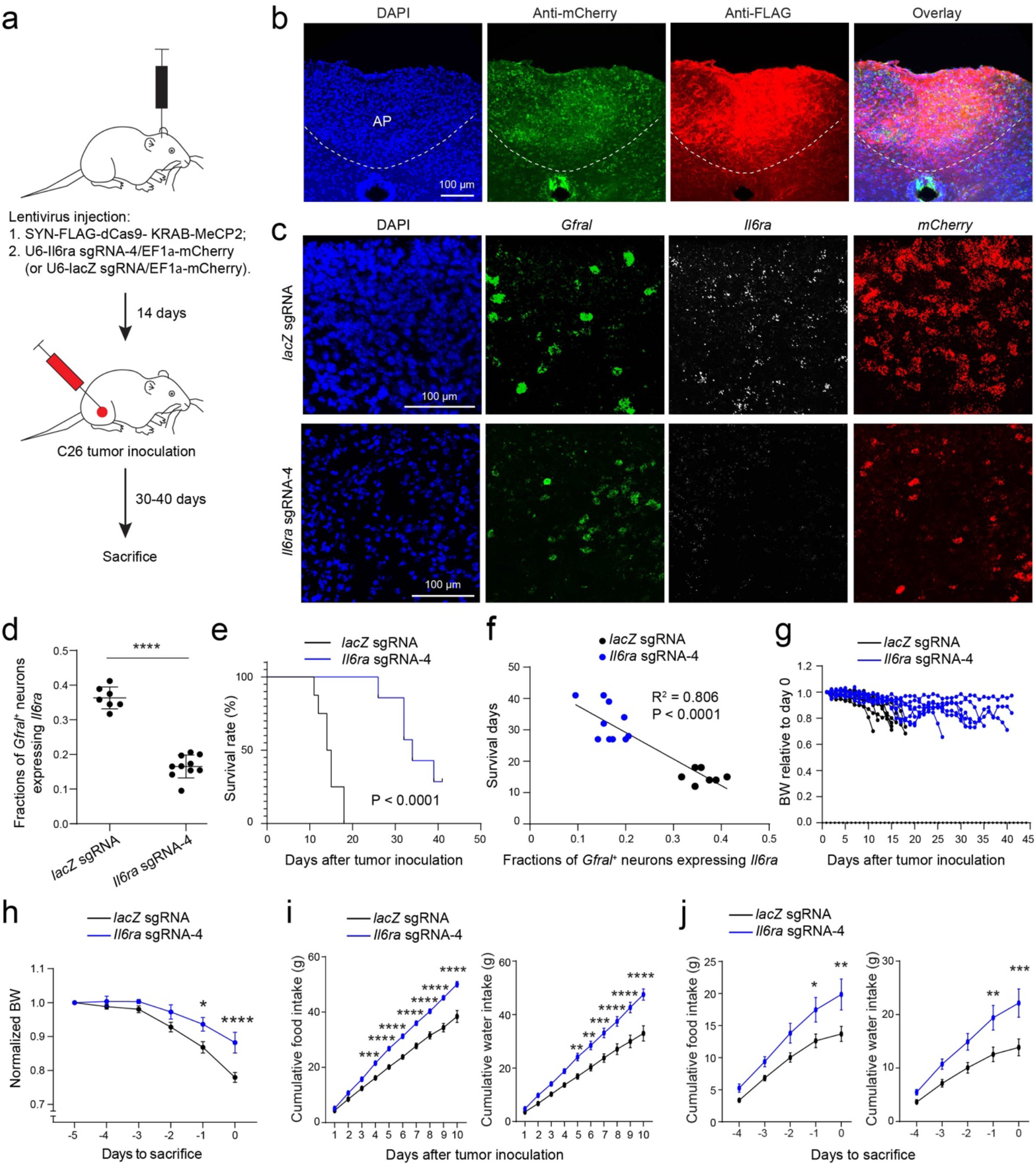
Suppression of *Il6ra* expression in AP neurons ameliorates cachexia in the C26 cancer model. **a**. A schematic of the experimental procedure. **b**. Confocal immunohistochemical images of a coronal brain section from a representative mouse, showing the infection of AP cells with lentiviruses expressing the sgRNA (tagged with mCherry) and dCas9-KRAB-MeCP2 (tagged with FLAG). mCherry and FLAG were recognized by antibodies. **c**. Confocal images showing the expression of *Gfral, Il6ra*, and *mCherry* in AP cells detected with smFISH. *mCherry* marks the cells infected with the CRISPR/dCas9 system containing the *lacZ* sgRNA (top) or *Il6ra* sgRNA-4 (bottom). **d**. Quantification of the fractions of *Gfral*^*+*^ neurons expressing *Il6ra* in the AP of individual mice (*lacZ* sgRNA group, n = 7, *Il6ra* sgRNA-4 group, n = 10; t = 12.42, ****P = 2.7 × 10^−9^, t test). **e**. Survival curves of the mice after tumor inoculation (*lacZ* sgRNA group, n = 8, *Il6ra* sgRNA-4 group, n = 7; P < 0.0001, Mantel-Cox test). **f**. Relationship between survival days and the fractions of *Gfral*^*+*^ neurons expressing *Il6ra* (n = 17 mice, R^2^ = 0.806, P < 0.0001 by a linear regression). **g**. Bodyweight of individual mice relative to their bodyweight on the day of tumor inoculation. **h**. Average bodyweight normalized to that on day - 5 (*lacZ* sgRNA group, n = 8, *Il6ra* sgRNA-4 group, n = 7; F(1,78) = 23.63, P = 9.6 × 10^−6^, *P = 0.012, ****P = 0.00004, two-way repeated-measures (RM) ANOVA with Sidak’s post hoc test). **i**. Cumulative food (left) and water (right) intake of the mice after tumor inoculation (food: F(1,130) = 253.6, P < 1 × 10^−15^; day 4, ***P = 0.0007, day 5, ****P = 9.2 × 10^−6^, day 6, ****P = 5.5 × 10^−7^, day 7, ****P = 3.6 × 10^−8^, day 8, ****P = 4.23 × 10^−9^, day 9, ****P = 9.21 × 10^−13^, day 10, ****P = 2.4 × 10^−14^; water: F(1,130) = 125.2, P < 1 × 10^−15^; day 5, **P = 0.009, day 6, **P = 0.0019, day 7, ***P = 0.00024, day 8, ****P = 0.00003, day 9, ****P = 2.65 × 10^−7^, day 10, ****P = 3.58 × 10^−9^; two-way RM ANOVA with Sidak’s post hoc test). **j**. Cumulative food (left) and water (right) intake of the mice in the 5 days before sacrifice (food: F(1, 65) = 24.83, P = 4.9 × 10^−6^; *P = 0.033, **P = 0.0034; water: F(1, 65) = 31.32, P = 4.7×10^−7^; **P = 0.0064, ***P = 0.0006; two-way RM ANOVA with Sidak’s post hoc test).

Compared with the *lacZ* sgRNA control group, the *Il6ra*-sgRNA-4 group had reduced *Il6ra* expression in AP neurons, including the *Gfral*^*+*^ neurons (Fig. 4c, d). Notably, the *Il6ra*-sgRNA-4 group had markedly increased lifespans (Fig. 4e), an effect that was inversely correlated with the percentage of *Gfral*^*+*^ neurons that had detectable *Il6ra* expression (Fig. 4f). The *Il6ra*-sgRNA-4 group also had reduced bodyweight loss (Fig. 4g, h), increased food and water intake (Fig. 4i, j), increased blood glucose levels (Extended Data Fig. 4a), and had a tendency to reduce tissue loss (Extended Data Fig. 4b, c). At the endpoint of the experiment, mice in the *Il6ra*-sgRNA-4 group had larger tumor and spleen compared with mice in the control group (Extended Data Fig. 4d, e), presumably because of the increase in lifespan in the former group.

In a separate experiment, the *Il6ra*-sgRNA-4 mice and the *lacZ*-sgRNA control mice were sacrificed on the same day, as soon as the latter had developed cachexia (Extended Data Fig. 5a). This design ensured that the two groups had tumor for the same duration and thus had comparable tumor and spleen mass (Extended Data Fig. 5g, h). Consistent with the above observations, the *Il6ra*-sgRNA-4 group had reduced bodyweight loss and tissue loss (Extended Data Fig. 5b-d), even though these mice had similar food and water intake to the control mice before the termination of this experiment (Extended Data Fig. 5e, f). Interestingly, the *Il6ra*-sgRNA-4 mice had reduced Fos expression compared with the *lacZ* sgRNA mice in the AP, PBN, and PVN (Extended Data Fig. 5i, j), suggesting that suppression of *Il6ra* expression in AP neurons lowers the hyperactivity in the AP network.

Additional experiments showed that the other sgRNA, the *Il6ra* sgRNA-6, had similar effects to those of *Il6ra* sgRNA-4 (Extended Data Fig. 6). Together, these results indicate that the IL-6Rα on AP neurons, especially that on *Gfral*^*+*^ neurons, is a critical mediator of IL-6 function in the development of cancer-associated cachexia.

### Inhibition of Gfral^+^ neurons ameliorates cancer-induced anorexia

Given the result that the survival of mice in the C26 model was correlated with the suppression of *Il6ra* in *Gfral*^*+*^ neurons in the AP (Fig. 4f), and previous findings that Gfral and its ligand GDF-15 (Growth/differentiation factor 15) are involved in cancer-associated cachexia^28,29^, we reasoned that the activity of *Gfral*^*+*^ AP neurons contributes to the development of this syndrome. To test this hypothesis, we sought to selectively manipulate these neurons using the *Gfral-p2a-Cre* mice^50^ in combination with a Cre-dependent viral approach. As the C26 model necessitates the use of Balb/c or CD2F1 mice^60^, it is incompatible with the *Gfral-p2a-Cre* mice which have a C57BL/6 genetic background. Therefore, for this experiment, we used the implantable Lewis lung carcinoma (LLC) model, another established murine cancer model exhibiting features of cachexia, albeit milder than those of the C26 model^18,23,27^.

We first measured the levels of IL-6 and GDF-15 in the blood at different timepoints of cancer progression in this model. Plasma IL-6 and GDF-15 were both increased at around two weeks following tumor inoculation (Fig. 5a). To specifically inhibit the activity of *Gfral*^*+*^ neurons, we injected the AP of *Gfral-p2a-Cre* mice bilaterally with an adeno-associated virus (AAV) expressing the tetanus toxin light chain (TeLC, which blocks neurotransmitter release^61^), or GFP (as a control) in a Cre-dependent manner. Two weeks later, these mice were inoculated with the LLC (Fig. 5b, c). Interestingly, compared with the GFP mice, the TeLC mice exhibited increased food intake at the late stage of cancer progression (Fig. 5d) and increased fat and muscle mass at the endpoint (Fig. 5e). Moreover, the TeLC mice showed reduced Fos expression than the GFP mice in the AP, PBN, PVN, CeA and BNST (Fig. 5f, g). The two groups had similar tumor and spleen mass (Fig. 5h). These results indicate that reducing *Gfral*^*+*^ AP neuron activity ameliorates the cachectic phenotypes. In addition, other areas in the AP network are likely also involved in this process.

**Figure 5.**
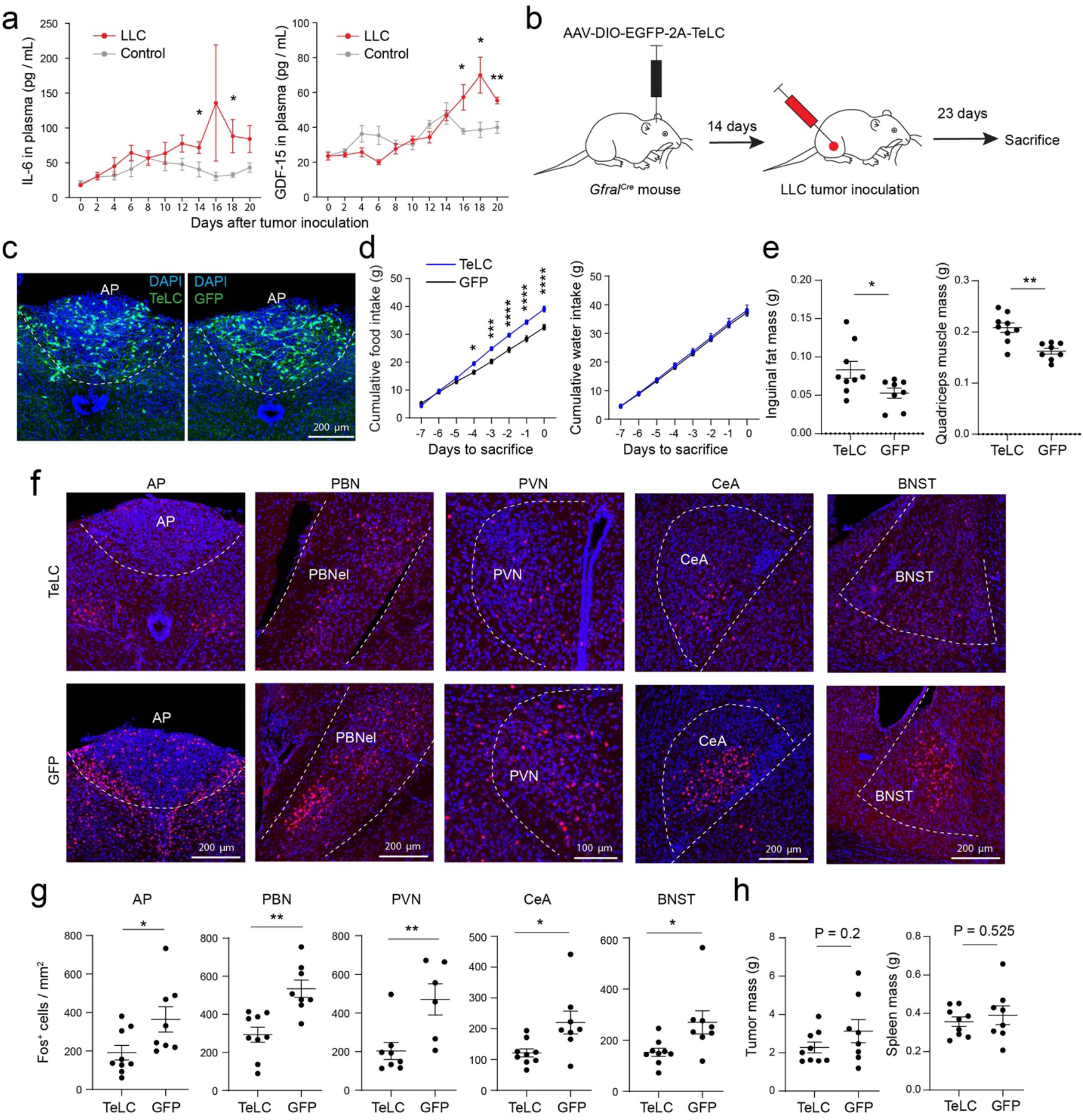
Inhibition of *Gfral*^*+*^ AP neurons ameliorates cachexia in the Lewis lung cancer (LLC) model. **a**. Plasma IL-6 (left) and GDF-15 (right) concentrations during cancer progression (IL-6: control, 7 mice at all timepoints except day 2 and day 8, where there are 5 mice; LLC, 8 mice at all timepoints except day 4 where there are 6 mice, day 6 and day 12 where there are 11 mice, and day 10 and day 18 where there are 7 mice; *P < 0.05; GDF-15: control, 5 mice at all timepoints except day 0 where there are 8 mice, and day 14 where there are 3 mice; LLC, 5 mice at all timepoints except day 0 where there are 7 mice; *P < 0.05, **P < 0.01; multiple unpaired t-tests at each timepoint with false discovery rate adjusted with the two-stage step-up method). **b**. A schematic of the experimental procedure. **c**. Confocal immunohistochemical images of coronal brain sections from two representative mice, showing the infection of *Gfral*^*+*^ AP neurons with an AAV expressing TeLC (left) or GFP only (right). **d**. Cumulative food (left) and water (right) intake of the mice in the 7 days before sacrifice (TeLC group, n = 9, GFP group, n = 8; food: F(1,120) = 78.04, P = 10^−14^; day -4, *P = 0.03, day -3, ***P = 0.0002, day -2, ****P = 0.00002, day -1, ****P = 5 × 10^−7^, day 0, ****P = 5.8 × 10^−8^; water: F(1,120) = 1.256, P = 0.26; two-way repeated-measures ANOVA with Sidak’s post hoc test. **e**. Inguinal fat (left) and quadriceps muscle (right) mass of the mice at 23 days after tumor inoculation (TeLC group, n = 9, GFP group, n = 8; fat: t = 2.3, *P = 0.0363; muscle: t = 4.04, **P = 0.001; t test). **f**. Confocal images showing Fos expression in different brain areas in the mice where *Gfral*^*+*^ AP neurons were infected with the AAV expressing TeLC (top) or GFP only (bottom). **g**. Quantification of Fos^+^ cells in different brain areas (TeLC group, n = 9 mice, GFP group, n = 8 mice; AP, t = 2.33, *P = 0.0343; PBN, t = 4.04, **P = 0.0011; PVN, t = 3.09, **P = 0.0094; CeA, t = 2.62, *P = 0.019; BNST, t = 2.55, *P = 0.022; unpaired t test). **h**. Tumor (left) and spleen (right) mass of the mice at 23 days after tumor inoculation (TeLC group, n = 9, GFP group, n = 8; tumor: t = 1.34, P = 0.2; spleen: t = 0.65, P = 0.525; t test).

## Discussion

Our study identifies AP neurons as a critical mediator of the function of IL-6 that leads to cancer-associated cachexia in mice. We show that, first, circulating IL-6 rapidly enters the AP and causes AP neuron activation within hours. Second, periphery tumor results in increased IL-6 and hyperactivity in the AP, and induces enhanced excitatory synaptic drive onto AP neurons. Third, neutralization of IL-6 in the brain of tumor-bearing mice, via i.c.v. infusion of an IL-6 antibody, prevents cachexia, counteracts the cachexia-associated hyperactivity in the AP network, and markedly prolongs lifespan. Fourth, specific suppression of *Il6ra* in AP neurons with CRISPR/dCas9 interference has similar effects. Lastly, specific suppression of the activities of *Gfral*^+^ neurons, which partially overlap with *Il6ra*^+^ cells in the AP (Fig. 1h) and are involved in the effects of *Il6ra* suppression (Fig. 4f), also ameliorates cancer-associated cachectic phenotypes and AP network hyperactivity.

The AP network showing cachexia-associated hyperactivity includes the PBN, the PVN, the BNST, and the CeA besides the AP. These structures are interconnected^50,53-57^ and have been implicated in regulating feeding behavior and metabolism^50,56,62-69^. In particular, the AP and the neighboring nucleus tractus solitarii (NTS), as well as the PBN and the PVN have previously been implicated in cancer-associated cachexia^17,27-29^. The AP sends direct projections to the PBN and the NTS, and the NTS also directly projects to the PBN as well as the PVN^50,57,70^. Given previous findings that both the PBN^27,50,55,56,70-72^ and the PVN^16,73,74^ are involved in feeding suppression, it is possible that the AP drives cancer-associated anorexia via the AP→PBN, the AP→NTS→PBN, or the AP→NTS→PVN pathway.

A notable observation is that the AP also drives weight loss independent of anorexia during cancer progression (Extended Data Fig. 5), consistent with findings that cancer-associated cachexia involves active catabolic processes in addition to anorexia, and the tissue wasting can only be partially reversed by nutritional support^18-20^. Interestingly, multiple nuclei in the AP network are connected with mechanisms that promote catabolic process in peripheral organs^62,63,66,69,75^, providing an anatomical basis for this function of the AP.

Recent studies indicate that Gfral is exclusively expressed by neurons in the AP and the NTS^76-79^, and systemic administration of GDF-15 activates GFRAL^+^ neurons in the AP and induces vomiting and anorexia^28,50,80-83^. Furthermore, neutralization of Gfral or GDF-15 with antibodies ameliorates cancer-associated cachectic phenotypes in animals^28,29^. Thus, GDF-15 may also influence cancer-associated cachexia, like IL-6, through the AP network. However, as GDF-15 functions as a central alert to the organism in response to a broad range of stressors^84^, including infection, blockade of GDF-15/GFRAL is likely to have detrimental effects if used as a therapeutic strategy. Indeed, it has recently been shown that GDF-15 is essential for surviving bacterial and viral infections^65^.

IL-6 has long been known as a key contributor to cancer-associated cachexia^18-20,23,24,35,36^. Efforts exploring IL-6 as a potential therapeutic target thus far have been focused on peripheral IL-6 or IL-6 receptors, and relied on systemic application of antibodies against these molecules^18,23,38^. However, such systemic approach may not be effective in reducing IL-6 signaling in the brain. Furthermore, as IL-6 is a pleiotropic cytokine essential for immune and metabolic functions, with receptors widely distributed in the entire organism^23,85^, systemic neutralization of IL-6 or its receptors will compromise these functions globally and likely cause severe side effects^86,87^. Our results suggest that targeting IL-6 signaling in the brain, or more specifically in the AP, could be an effective treatment for cancer-associated cachexia.

## Methods

### Mice

Male mice aged 2–4 months were used in all the experiments. Mice were housed under a 12-h light/dark cycle (7 a.m. to 7 p.m. light) in groups of 2–5 animals, with a room temperature (RT) of 22 °C and humidity of 50%. Food and water were available *ad libitum* before experiments. All experimental procedures were approved by the Institutional Animal Care and Use Committee of Cold Spring Harbor Laboratory and performed in accordance with the US National Institutes of Health guidelines.

The Balbc mice (strain number: 000651) were purchased from Jackson laboratory. The *Gfral-p2a-Cre* mice was generated by Stephen Liberles^50^.

### Immunohistochemistry

Immunohistochemistry experiments were performed following standard procedures. Briefly, mice were anesthetized with Euthasol (0.2 ml; Virbac, Fort Worth, Texas, USA) and transcardially perfused with 30 ml of PBS, followed by 30 ml of 4% paraformaldehyde (PFA) in PBS. Brains were extracted and further fixed in 4% PFA overnight followed by cryoprotection in a 30% PBS-buffered sucrose solution for 36 h at 4 °C. Coronal sections (50 μm in thickness) were cut using a freezing microtome (Leica SM 2010R). Brain sections were first washed in PBS (3 × 5 min), incubated in PBST (0.3% Triton X-100 in PBS) for 30 min at RT and then washed with PBS (3 × 5 min). Next, sections were blocked in 5% normal goat serum in PBST for 30 min at RT and then incubated with the primary antibody overnight at 4 °C. Sections were washed with PBS (5 × 15 min) and incubated with the fluorescent secondary antibody at RT for 2 h. After washing with PBS (5 × 15 min), sections were mounted onto slides with Fluoromount-G (eBioscience, San Diego, California, USA). Images were taken using an LSM 780 laser-scanning confocal microscope (Carl Zeiss, Oberkochen, Germany).

The primary antibodies and dilutions used in this study were: rabbit anti-Fos (1:500, Santa Cruz, sc-52), mouse anti-FLAG (1:1000, Sigma-Aldrich, F1804), rabbit anti-mCherry (1:1,000; Abcam, ab167453, GR3213077-3). The fluorophore-conjugated secondary antibodies and dilutions used were Alexa Fluor 488 goat anti-rabbit IgG (H + L; 1:500; A-11008, Invitrogen), Alexa Fluor 647 goat anti-rabbit IgG (H + L; 1:500; A-21244, Invitrogen), Alexa Fluor 594 goat anti-mouse IgG (H + L; 1:500; A-11005, Invitrogen).

### Retro-orbital injection of exogenous IL-6 and its detection in the brain

Biotinylated human IL-6 solution (Acrobiosystems, IL-6-H8218; 2 µg/ml dissolved in saline) was injected into Balbc mice (100 µl per mouse) via retro-orbital injection. For retro-orbital injection, briefly, the animal was anaesthetized with 2% isoflurane. A 27-gauge needle on a 0.5 mL insulin syringe was used for the injection. The animal was placed on its side on a heat pad. The gauge needle was inserted at approximately a 30-45° angle to the eye, lateral to the medial canthus, through the conjunctival membrane. There is a bit of resistance that causes the eye to retreat back into the sinus until the needle pierces through the conjunctiva. The needle was positioned behind the globe of the eye in the retro-bulbar sinus. The biotinylated human IL-6 solution was injected slowly into the retro-bulbar sinus. After the injection, the needle was removed gently and the animal was returned to homecage for recovery. 3 hours after the injection, the animals were sacrificed and transcardially perfused with 30 ml of PBS, followed by 30 ml of 4% paraformaldehyde (PFA) in PBS. Brains were extracted and further fixed in 4% PFA overnight followed by cryoprotection in a 30% PBS-buffered sucrose solution for 36 h at 4 °C. Coronal sections (50 μm in thickness) were cut using a freezing microtome (Leica SM 2010R). Brain sections were incubated in Streptavidin solution (1:1000, ThermoFisher, Alexa Fluor™ 647 conjugate, dissolved in 0.3% PBST) in room temperature for 2 hours. After washing with PBS (5 × 15 min), sections were mounted onto slides with Fluoromount-G (eBioscience). Images were taken using an LSM 780 laser-scanning confocal microscope (Carl Zeiss).

### Fluorescence *in situ* hybridization

Single molecule fluorescent *in situ* hybridization (smFISH) (RNAscope, ACDBio) was used to detect the expression of *Glp1r, Il6ra, Fos*, and *Gfral* mRNAs in the area postrema of Balbc mice. For tissue preparation, mice were first anesthetized under isoflurane and then decapitated. Their brain tissue was first embedded in cryomolds (Sakura Finetek, Catalog number 4566) filled with M-1 Embedding Matrix (Thermo Scientific, Catalog number 1310) then quickly fresh-frozen on dry ice. The tissue was stored at −80 °C until it was sectioned with a cryostat. Cryostat-cut sections (16-μm thick) containing the entire area postrema were collected through the rostro-caudal axis in a series of four slides, and quickly stored at −80 °C until processed. Hybridization was carried out using the RNAscope kit (ACDBio). On the day of the experiment, frozen sections were postfixed in 4% PFA in RNA-free PBS (hereafter referred to as PBS) at RT for 15 min, then washed in PBS, dehydrated using increasing concentrations of ethanol in water (50%, once; 70%, once; 100%, twice; 5 min each). Sections were then dried at RT and incubated with Protease IV for 30 min at RT. Sections were washed in PBS three times (5 min each) at RT, then hybridized. Probes against *Glp1r* (Catalog number 418851-C3, dilution 1:50), *Il6ra* (Catalog number 438931-O1, dilution 1:50), *Fos* (Catalog number 316921-C2, dilution 1:50), and *Gfral* (Catalog number 417021-C2, dilution 1:50) were applied to the area postrema sections. Hybridization was carried out for 2 h at 40 °C. After that, sections were washed twice in 1× Wash Buffer (Catalog number 310091; 2 min each) at RT, then incubated with the amplification reagents for three consecutive rounds (30 min, 15 min and 30 min, at 40 °C). After each amplification step, sections were washed twice in 1× Wash Buffer (2 min each) at RT. Finally, fluorescence detection was carried out for 15 min at 40 °C. Sections were then washed twice in 1× Wash Buffer (2 min each), incubated with DAPI for 2 min, washed twice in 1× Wash Buffer (2 min each), then mounted with a coverslip using mounting medium. Images were acquired using an LSM780 confocal microscope equipped with 20x and 40x lenses, and visualized and processed using ImageJ and Adobe Illustrator. Cell counting and mean fluorescence intensity quantification of the images were performed with ImageJ.

### Single guide RNA (sgRNA) design and lentiviral production for CRISPR/dCas9 interference

sgRNAs targeting the *Il6ra* transcription start site (TSS) were designed using CHOPCHOP^88^. Seven *Il6ra* sgRNAs (sgRNA-1 to sgRNA-7) as well as a sgRNA targeting the *lacZ* promoter (*LacZ* sgRNA) were cloned into the Lenti U6-sgRNA/Ef1a-mCherry plasmid (Addgene #114199), as described previously^89,90^. The eight sgRNA plasmids, Lenti SYN-FLAG-dCas9-KRAB-MeCP2 plasmid (Addgene #155365), and the two helper plasmids pCMV-VSV-G (Addgene #8454) and psPAX2 (Addgene #12260) were purified with the NucleoBond Xtra Midi EF kit (Takara 740420). *Il6ra* knockdown efficiency was assessed by transient transfection of sgRNA and dCas9-KRAB-MeCP2 into the mHypoA hypothalamic neural cell line (Cedarlane Labs, clone clu-175). 3 µg of each sgRNA plasmid and 3 µg of dCas9-KRAB-MeCP2 plasmid were co-nucleofected into 1 × 10^6^ mHypoA cells resuspended in a 1:1 mixture of Ingenio Electroporation reagent (Mirus Bio 50111) and OptiMEM (Gibco 31985062), using program A-033 on the Nucleofector 2b (Lonza). The cells were harvested 60 hours post-nucleofection. DAPI- and mCherry-positive cells were collected by FACS. The *Il6ra* mRNA was extracted and the knockdown efficiency was measured by RT-qPCR. The two most effective sgRNAs, sgRNA-4 (−23 to −41 of TSS) and sgRNA-6 (−163 to −182 of TSS), resulting in 67% and 35% knockdown of *Il6ra* expression in mHypoA cells, respectively, were used for *in vivo* experiments. FLAG-dCas9-KRAB-MeCP2, *Il6ra* sgRNA-4, *Il6ra* sgRNA-6, and *lacZ* sgRNA lentiviruses were produced in HEK293T cells. Lentiviral pellets were resuspended in 30 uL DPBS, aliquoted and flash-frozen on dry ice, and stored at −80°C. Physical and functional titers were obtained using the Lenti-X qRT-PCR Titration Kit (Takara 631235) and qPCR of genomic DNA following HEK293T transduction^91^, respectively.

| Lentiviral preparation | Physical titer<br>(genomic copies/uL) | Functional titer<br>(transducing units/uL) |
| --- | --- | --- |
| FLAG-dCas9-KRAB-MeCP2 (prep 1) | 3.0E+11 | 1.2E+07 |
| FLAG-dCas9-KRAB-MeCP2 (prep 2) | 8.0E+10 | 1.1E+07 |
| *Il6ra* sgRNA-4 | 3.0E+12 | 1.4E+08 |
| *Il6ra* sgRNA-6 | 5.0E+11 | 3.8E+07 |
| *lacZ* sgRNA | 3.0E+12 | 8.8E+07 |

#### Plasmids for lentiviral production

lenti SYN-FLAG-dCas9-KRAB-MeCP2 was a gift from Jeremy Day (Addgene plasmid # 155365; http://n2t.net/addgene:155365; RRID:Addgene_155365)

lenti U6-sgRNA/EF1a-mCherry was a gift from Jeremy Day (Addgene plasmid # 114199; http://n2t.net/addgene:114199; RRID:Addgene_114199)

pCMV-VSV-G was a gift from Bob Weinberg (Addgene plasmid # 8454; http://n2t.net/addgene:8454; RRID:Addgene_8454)

psPAX2 was a gift from Didier Trono (Addgene plasmid # 12260; http://n2t.net/addgene:12260; RRID:Addgene_12260)

### Adeno-associated viruses (AAVs)

The AAV-CMV-DIO-EGFP-2A-TeLC vector was a gift from Dr. Wei Xu at UT Southwestern. A custom virus (AAV-DJ) based on this vector was produced by WZ Biosciences Inc (Rockville, MD 20855). pAAV-hSyn-DIO-EGFP was purchased from Addgene (Watertown, MA 02472, USA). All viruses were aliquoted and stored at −80 °C until use.

### Stereotaxic surgery

All surgery was performed under aseptic conditions and body temperature was maintained with a heating pad. Standard surgical procedures were used for stereotaxic injection. Briefly, mice were anesthetized with isoflurane (3% at the beginning and 1% for the rest of the surgical procedure), and were positioned in a stereotaxic injection frame and on top of a heating pad maintained at 35 °C. A digital mouse brain atlas was linked to the injection frame to guide the identification and targeting of different brain areas (Angle Two Stereotaxic System, http://myNeuroLab.com). We used the following coordinates for injections in the area postrema: −7.65 mm from bregma, 0 mm lateral from the midline, and 4.7 mm vertical from the skull surface. For virus injection, we made a small cranial window (1–2 mm^2^) for each mouse, through which a glass micropipette (tip diameter, ∼5 μm) containing viral solution was lowered down to the target. For AAVs, about 0.3 μl of viral solution was injected. For lentiviruses, 0.2-0.3 μl of viral mixture (the dCas9 and the sgRNA viruses were mixed at a volume:volume ratio of 2:1) was injected. Viral solutions were delivered with pressure applications (5–20 psi, 5–20 ms at 1 Hz) controlled by a Picospritzer III (General Valve) and a pulse generator (Agilent). The speed of injection was ∼0.1 μl/10 min. We waited for at least 10 min following the injection before slowly removing the injection pipette. After injection, the incision was sealed by surgical sutures and the animal was returned to homecage for recovery.

### Colon-26 (C26) adenocarcinoma cells

C26 cells were cultured in complete growth medium consisting of RPMI 1640 medium with Glutamine (#11-875-093; Thermo Fisher) containing 10% of heat-inactivated Fetal Bovine Serum (FBS) (#10-438-026; Thermo Fisher) and 1x Penicillin-Streptomycin solution (#15-140-122; Thermo Fisher) under sterile conditions. 1x Trypsin-EDTA (#15400054; Thermo Fisher) was used for cell dissociation. Cells were resuspended in FBS-free RPMI and viable cells were counted using a Vi-Cell counter prior to subcutaneous injection of 2 × 10^6^ viable cells diluted in 100 μL RPMI into the right flank of each BALB/c mouse.

### Lewis lung carcinoma (LLC) cells

LL/2 (LLC1) cells were obtained from ATCC (American Type Culture Collection; #CRL-1642) and cultured in Dulbecco’s Modified Eagle’s Medium (DMEM) (#30-2002; ATCC) complete growth medium, with 10% of heat-inactivated FBS (#10-438-026; Thermo Fisher) and 1x Penicillin-Streptomycin solution (#15-140-122; Thermo Fisher) under sterile conditions. 1x Trypsin-EDTA (#15400054; Thermo Fisher) was used for cell dissociation. Cells were resuspended in FBS-free DMEM and viable cells were counted using a Vi-Cell counter prior to subcutaneous injection of. 2 × 10^6^ viable cells were diluted in 100 μL FBS-free DMEM and were subcutaneous injected into the right flank of each C57BL/6 mouse.

### Intracerebroventricular (i.c.v.) infusion of anti-IL-6

On day 10 or 12 post-C26 injection, an osmotic device of 200 uL volume and a release rate of 0.5 uL/hour consisting of a cannula, connecting line, metal flow moderator and pump (#AP-2001; Alzet) was placed in a subcutaneous pocket and stereotactically implanted into the right lateral ventricle of the C26-tumour bearing BALB/c mice for a period of 14 days. Prior to use, the infusion device was assembled and equilibrated in saline overnight. The pump was filled with either the InVivoMAb rat anti-mouse IL-6 (clone MP5-20F3, #BE0046; BioXCell) or an InVivoMAb rat IgG1 isotype control (anti-HRPN, #BE0088; BioXCell). Both antibodies were diluted in PBS to achieve continuous infusion of a 5 mg/mL dose. Pump replacement surgery was performed after 14 days. The coordinate for targeting the lateral ventricle was −0.5 mm from bregma, 1.25 mm lateral from the midline, and 2.5 mm vertical from the skull surface.

### Measuring bodyweight, food intake, and water intake

Food and water intake monitoring cages (BioDAQ Unplugged, Research Diets, Inc., New Brunswick, NJ 08901 USA) were used to measure the food intake and water intake of the animals. Mice were singly housed in these cages. Food and water were placed in an extended hopper which can be reached by the animal. The bodyweight of the animal, weight of the food and water in the hopper were measured daily at 4 pm. The cachectic mice which lost >20% of bodyweight were sacrificed and the tissues were collected for further analysis. Blood glucose concentrations were measured from whole venous blood using an automatic glucose monitor (Bayer HealthCare Ascensia Contour).

### Blood and plasma measurements

Blood glucose concentrations were measured from whole venous blood using an automatic glucose monitor (Bayer HealthCare Ascensia Contour). Tail vein bleeding was performed using a scalpel via tail venesection without restraint. Blood samples were collected from tail bleed using heparin-coated hematocrit capillary tubes to avoid coagulation. Samples were then centrifuged at 14,000 rpm for 5 min at 4°C. Plasma was collected in a new tube, snap frozen in liquid nitrogen and stored at −80°C. IL-6 and GDF-15 levels were measured in plasma using the mouse IL-6 Quantikine ELISA Kit (#M6000B; R&D Systems) and the Mouse/Rat GDF-15 Quantikine ELISA Kit (#MGD150; R&D) respectively.

### Brain tissue lysis and IL-6 quantification

Mice were transcardially perfused with saline and the area postrema was collected, snap frozen in liquid nitrogen and stored at −80°C until further analysis. Tissue was placed into 2-mL round-bottom homogenizer tubes pre-loaded with Stainless Steel beads (#69989; Qiagen) and filled up with lysis buffer (#AA-LYS-16ml; RayBiotech) supplemented with Protease Inhibitor Cocktail (#AA-PI; Raybiotech) and Phosphatase Inhibitor Cocktail Set I (#AA-PHI-I; RayBiotech). Samples were homogenized in Tissue Lyser II (#85300; Qiagen) for 5 minutes and then lysates were centrifuged at 4°C for 20 minutes at maximum speed. The supernatant was harvested and kept on ice if testing fresh or sored at −80°C. The Bicinchoninic Acid (BCA) Method was used to determine protein concentration in lysates. IL-6 levels were quantified in the lysates using a Mouse IL-6 ELISA specific for lysates (#ELM-IL6-CL-1; RayBiotech).

### *In vitro* electrophysiology

Acute slices were obtained from two-to three-month-old mice. Mice were anaesthetized with isoflurane (4%) before rapid decapitation. The brain was rapidly removed, and coronal slices (300 μm) containing the AP were cut using a HM650 Vibrating-blade Microtome (Thermo Fisher Scientific). Slices were cut in ice-cold dissection buffer (110.0 mM Choline chloride, 25.0 mM NaHCO_3_, 1.25 mM NaH_2_PO_4_, 2.5 mM KCl, 25.0 mM glucose, 0.5 mM CaCl_2_, 7.0 mM MgCl_2_, 11.6 mM ascorbic acid, and 3.1 mM pyruvic acid, and bubbled with 95% O_2_ and 5% CO_2_) and subsequently transferred to a recovery chamber containing artificial cerebrospinal fluid (ACSF) solution (containing 118 mM NaCl, 2.5 mM KCl, 26.2 mM NaHCO_3_, 1 mM NaH_2_PO_4_, 20 mM Glucose, 2 mM CaCl_2_ and 2 mM MgCl_2_, pH 7.4, and saturated with 95% O_2_ and 5% CO_2_) at 34°C. The slices were maintained at 34°C for at least 40 minutes and subsequently at room temperature (20-24°C). Recordings were made in a continuously flow of ACSF and bubbled with 95% O_2_/5% CO_2_.

Whole-cell patch-clamp recordings were obtained with Multiclamp 700B amplifiers and pCLAMP 10 software (Molecular Devices; Sunnyvale, California, USA) and was guided using an Olympus BX51 Microscope equipped with both transmitted and epifluorescence light sources (Olympus Corporation, Shinjuku, Tokyo, Japan).

Synaptic responses were recorded at holding potentials of −70mV (for AMPA receptor-mediated responses), and 0 mV (for GABAA receptor-mediated responses) and were low-pass filtered at 1 kHz. The internal solution for voltage-clamp experiments contained 115 mM Cesium methanesulfonate, 20 mM CsCl, 10 mM HEPES, 2.5 mM MgCl_2_, 4 mM Na_2_-ATP, 0.4 mM Na_3_-GTP, 10 mM Na-phosphocreatine, and 0.6 mM EGTA, pH 7.2. Miniature EPSCs were recorded in the presence of tetrodotoxin (1 µM) and picrotoxin (100 µM). Spontaneous IPSCs were recorded in the presence of AP-5 (100 µM) and CNQX (5 µM). The EPSCs and IPSCs were analyzed using Mini Analysis software (Synaptosoft).

### Statistical Analysis

All statistics are indicated where used. Statistical analyses were conducted using GraphPad Prism version 6.0 (GraphPad Software, Inc., La Jolla, CA). Statistical comparisons were performed using Student’s t test or ANOVA. All comparisons were two tailed. Statistic hypothesis testing was conducted at a significance level of 0.05.

## Data availability

The authors declare that the data supporting the findings of this study are available within the paper and its supplementary information files.

## Acknowledgements

We thank Dr. Stephen D. Liberles for sharing the *Gfral-p2a-Cre* mice. We thank Dr. Roberto Malinow and Dr. Chuchu Zhang for comments on earlier versions of the manuscript. We thank Radhashree Sharma for technical assistance and members of the Li laboratory for helpful discussions. This work was supported by The Mark Foundation for Cancer Research (33300111, T.J.), developmental funds from CSHL Cancer Center Support Grant (5P30CA045508, T.J.), National Institutes of Health (NIH) (R01MH101214, R01MH108924, R01NS104944, R01DA050374, B.L.), and the Cold Spring Harbor Laboratory and Northwell Health Affiliation (B.L.).

## Author Contributions

Q.S., D.V.D.L., M.F., B.G, M.W., J.T., T.J. and B.L. designed research; Q.S., D.V.D.L., M.F., B.G, and M.W. performed research; Q.S., D.V.D.L., M.F., B.G, M.W., J.T., T.J. and B.L. analyzed data; and Q.S. and B.L. wrote the paper with inputs from all authors.

## FIGURES and SUPPLEMENTARY FIGURES

**Extended Data Figure 1.**
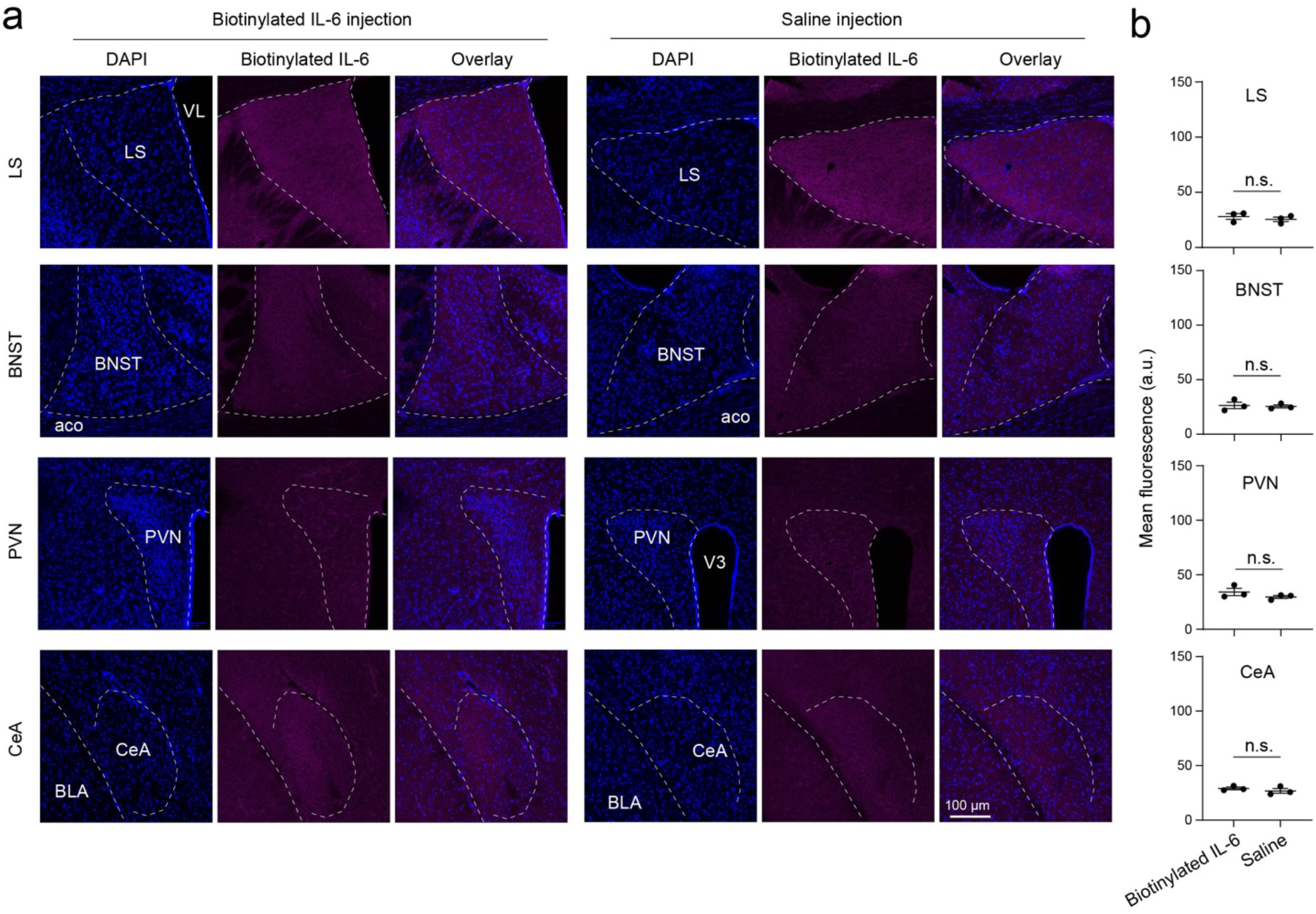
Circulating IL-6 does not reach brain areas other than the AP. **a**. Confocal images showing the lack of exogenous IL-6 signals in different brain areas of mice received biotinylated IL-6 (left) compared with mice received saline (right) via retro-orbital injection. **b**. Quantification of the fluorescence signals from fluorescence-conjugated avidin in different brain areas, which recognizes the biotinylated exogenous IL-6 (n = 3 mice for each group; LS, t = 0.765, P = 0.487; BNST, t = 0.282, P = 0.792; PVN, t = 1.277, P = 0.271; CeA, t = 0.892, P = 0.423; t test).

**Extended Data Figure 2.**
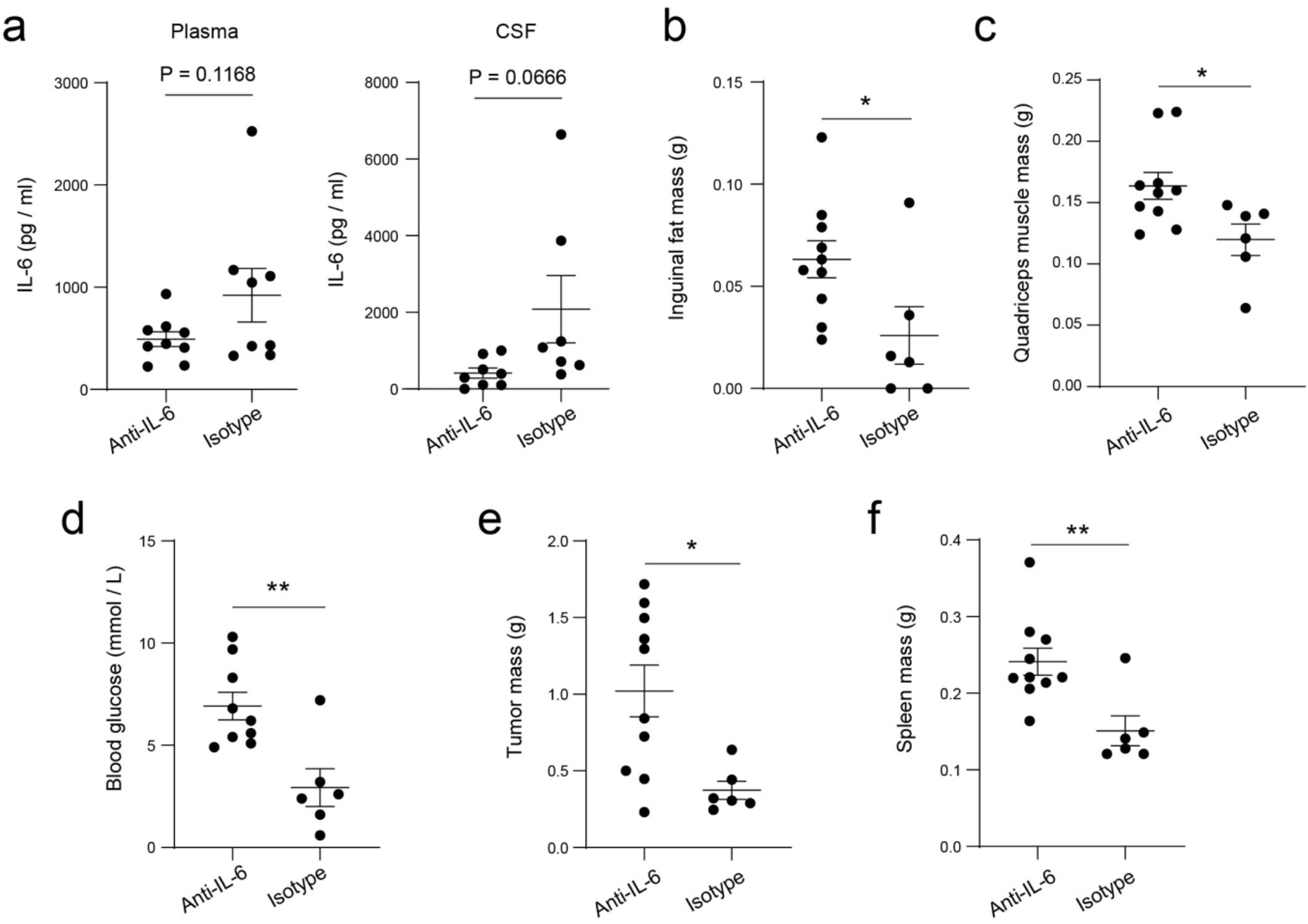
Intracerebroventricular (i.c.v.) infusion of IL-6 antibody improves the physiological conditions of mice despite tumor growth in the C26 cancer model. **a**. IL-6 levels in the plasma (left) and cerebrospinal fluid (CSF; right). Plasma IL-6: anti-IL-6 group, n = 9 mice, isotype control antibody group, n = 8 mice; t = 1.664, P = 0.1168; CSF IL-6: anti-IL-6 group, n = 8 mice, isotype control antibody group, n = 7 mice; t = 2.002, P = 0.0666; t test. **b**. Inguinal fat mass (anti-IL-6 group, n = 10, isotype control antibody group, n = 6; t = 2.334, *P = 0.035, t test). **c**. Quadriceps muscle mass (anti-IL-6 group, n = 10, isotype control antibody group, n = 6; t = 2.538, *P = 0.0237, t test). **d**. Blood glucose levels at the endpoint (anti-IL-6 group, n = 9 mice, isotype control antibody group, n = 6 mice; t = 3.555, **P = 0.0035, t test). **e**. Tumor mass (anti-IL-6 group, n = 10, isotype control antibody group, n = 6; t = 2.864, *P = 0.0125, t test). **f**. Spleen mass (anti-IL-6 group, n = 10, isotype control antibody group, n = 6; t = 3.274, **P = 0.0055, t test).

**Extended Data Figure 3.**
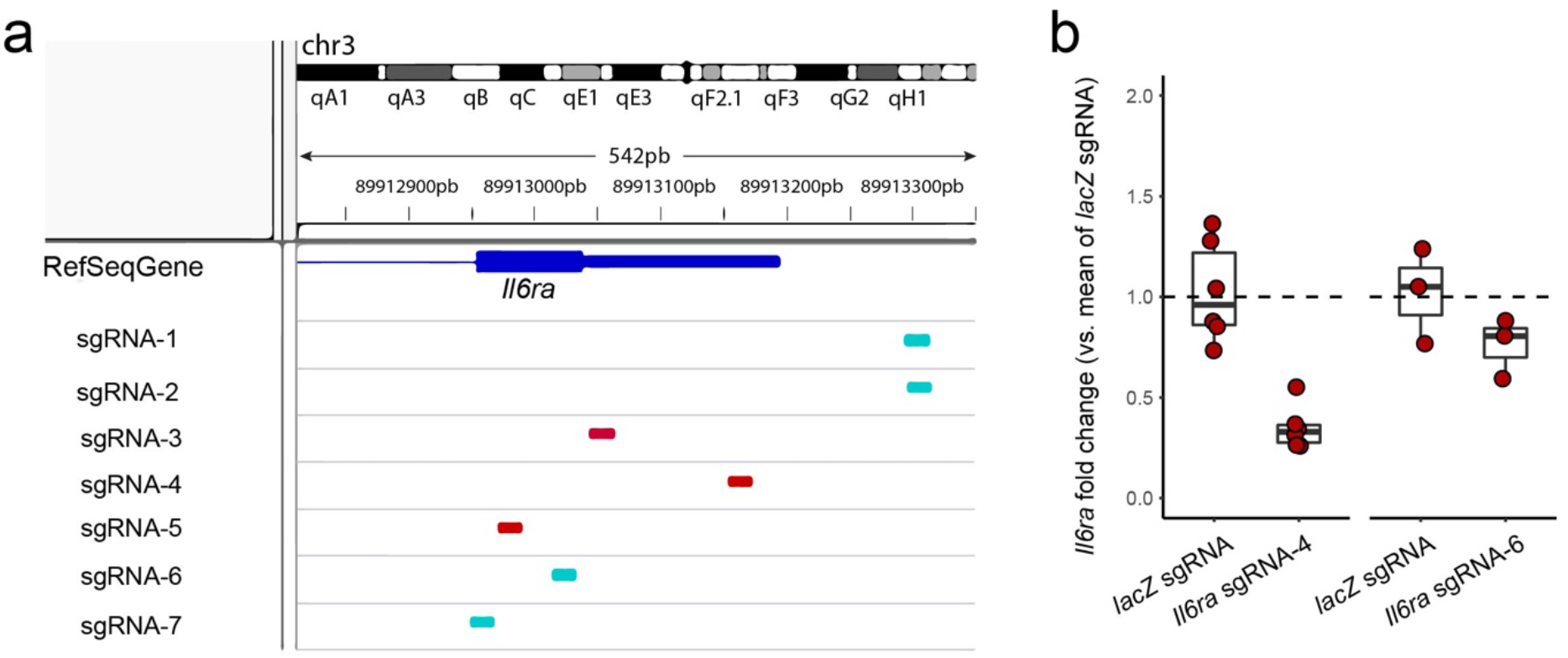
Design and characterization of sgRNAs for the CRISPR/dCas9 system to suppress *Il6ra* expression. **a**. Transcription start site (TSS) targeting positions of the different sgRNAs. Cyan, template strand; red, non-template strand. **b**. *In vitro* characterization of the efficacy of the CRISPR/dCas9 system with different sgRNAs. Plasmids expressing dCas9-KRAB-MeCP2 and each of the sgRNAs were co-transfected into mHypoA cell line. 60 hours after the transfection, the expression of *Il6ra* in these cells was measured by qPCR (*Il6ra* sgRNA-4, n = 6 plates, *lacZ* sgRNA, n = 6 plates, P = 0.0005785; *Il6ra* sgRNA-6, n = 3 plates, *lacZ* sgRNA, n = 3 plates, P = 0.1965; t test).

**Extended Data Figure 4.**
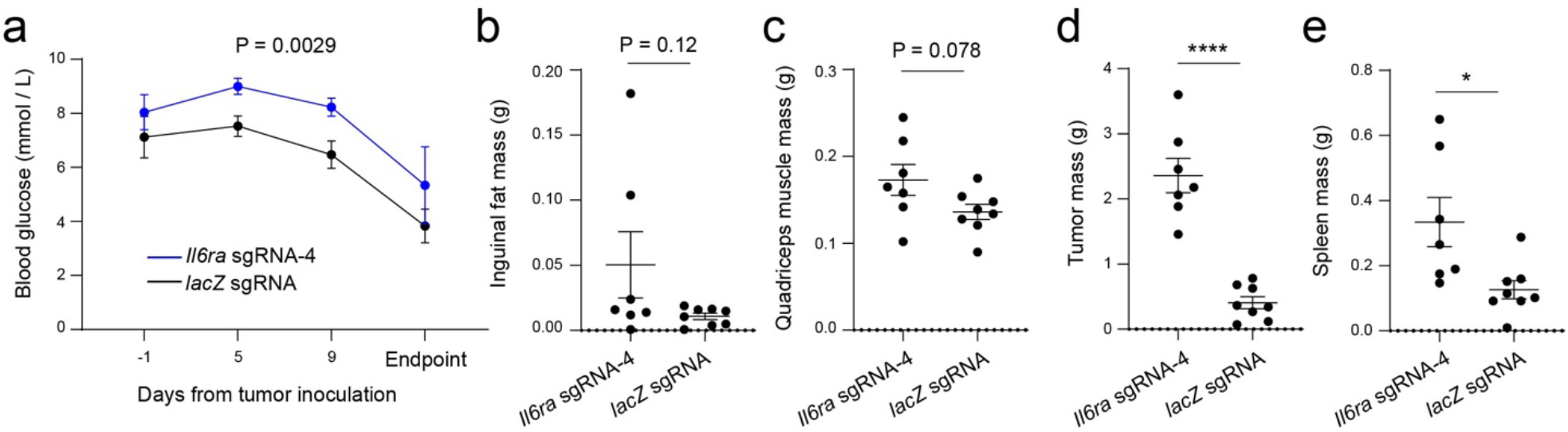
Suppression of *Il6ra* expression in AP neurons improves the physiological conditions of mice despite tumor growth in the C26 cancer model. *Il6ra* sgRNA-4 group, n = 7 mice, *lacZ* sgRNA group, n = 8 mice. **a**. Blood glucose levels at different time points (F(1,49) = 9.834, P = 0.0029; two-way repeated-measures ANOVA with Sidak’s post hoc test). **b**. Inguinal fat mass (t = 1.657, P = 0.12, t test). **c**. Quadriceps muscle mass (t = 1.92, P = 0.078, t test). **d**. Tumor mass (t = 7.32, ****P = 5.86 × 10^−6^, t test). **e**. Spleen mass (t = 2.71, P = 0.018, t test).

**Extended Data Figure 5.**
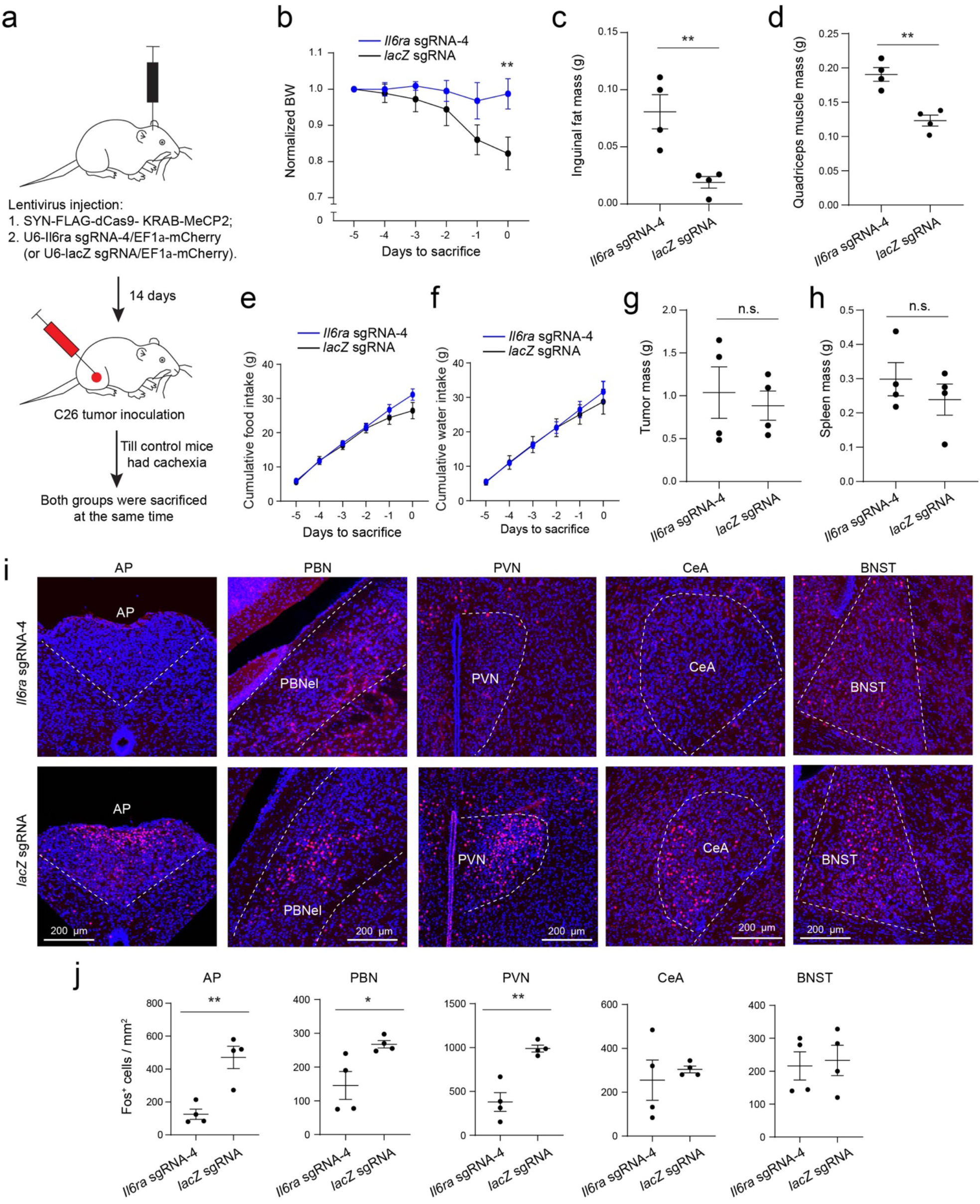
Suppression of *Il6ra* expression in AP neurons ameliorates cachexia and reduces the hyperactivity in the AP network in the C26 cancer model. *lacZ* sgRNA group, n = 4 mice, *Il6ra* sgRNA-4 group, n = 4 mice. **a**. A schematic of the experimental procedure. When one animal in the *lacZ* sgRNA (control) group became cachectic, that animal and a randomly selected animal in the *Il6ra* sgRNA-4 group were sacrificed to check Fos expression and other phenotypes. **b**. Average bodyweight normalized to that on day -5 (F(1,36) = 10.45, P = 0.0026, **P = 0.0072, two-way repeated-measures (RM) ANOVA with Sidak’s post hoc test). **c**. Inguinal fat mass (t = 3.941, **P = 0.0079, t test). **d**. Quadriceps muscle mass (t = 5.26, **P = 0.0019, t test). **e, f**. Cumulative food (e) and water (f) intake of the mice in the 5 days before sacrifice (food: F(1, 36) = 3.239, P = 0.0803; water: F(1, 36) = 0.3045, P = 0.5845. two-way RM ANOVA with Sidak’s post hoc test). **g, h**. Tumor (g) and spleen (h) mass of the mice (tumor: t = 0.444, P = 0.6725; spleen: t = 0.898, P = 0.404; t test). **i**. Confocal images showing Fos expression in different brain areas in representative mice of the two groups. **j**. Quantification of Fos^+^ cells in different brain areas (AP, t = 4.617, **P = 0.0036; PBN, t = 2.849, *P = 0.029; PVN, t = 5.332, **P = 0.0018; CeA, t = 0.5267, P = 0.617; BNST, t = 0.27, P = 0.796; unpaired t test).

**Extended Data Figure 6.**
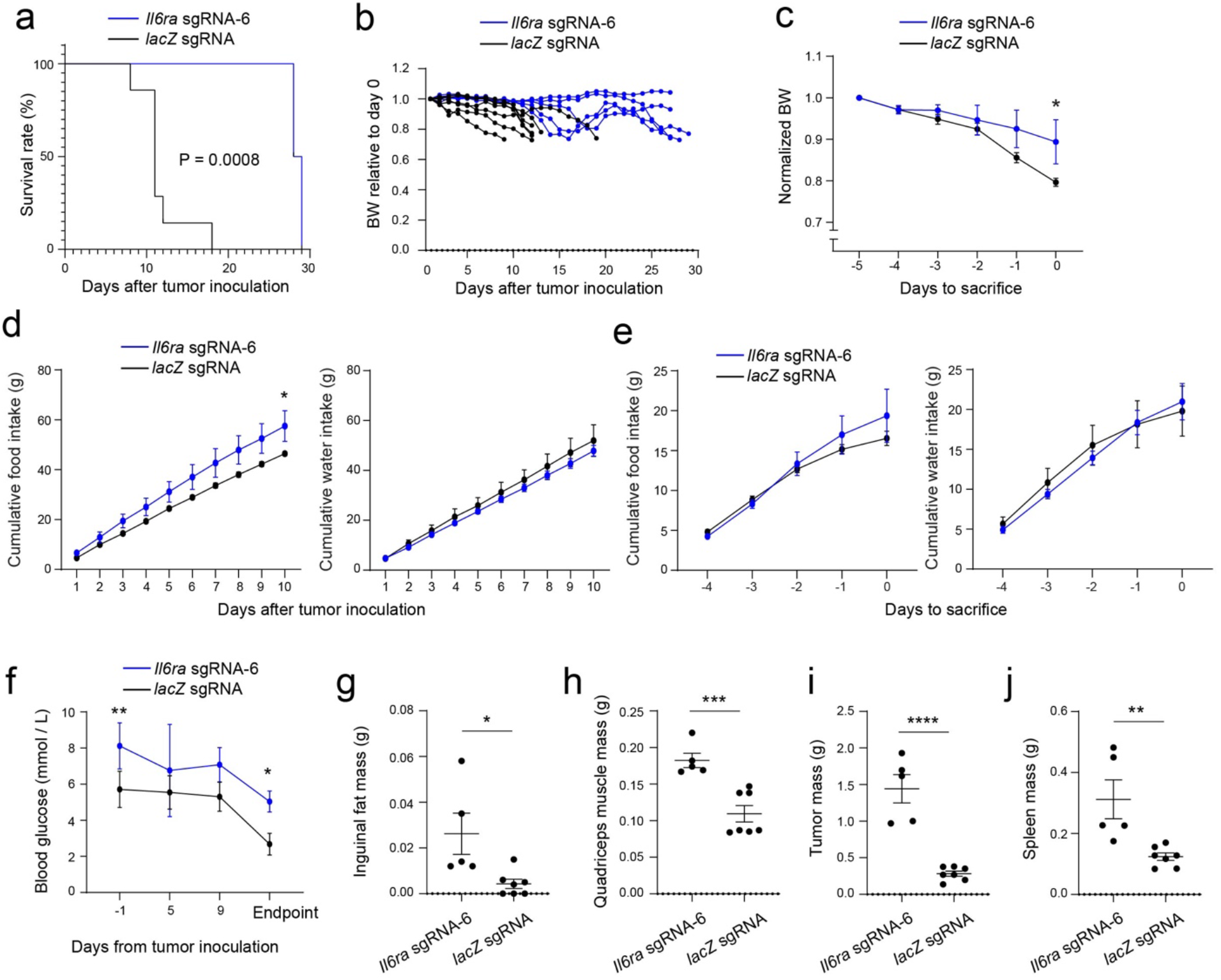
Suppression of *Il6ra* expression in AP neurons ameliorates cachexia in the C26 cancer model. *Il6ra* sgRNA-6 group, n = 5 mice, *lacZ* sgRNA group, n = 7 mice. **a**. Survival curves of the mice after tumor inoculation (P = 0.0008, Mantel-Cox test). **b**. Bodyweight of individual mice relative to their bodyweight on the day of tumor inoculation. **c**. Average of bodyweight normalized to that on day -5 (F(1,60) = 8.287, P = 0.0055; day 0, *P = 0.011, two-way repeated-measures (RM) ANOVA with Sidak’s post hoc test). **d**. Cumulative food (left) and water (right) intake of the mice after tumor inoculation (food: F(1,100) = 33.61, P = 7.89 × 10^−8^, *P = 0.049; water: F(1,100) = 1.977, P = 0.163; two-way RM ANOVA with Sidak’s post hoc test). **e**. Cumulative food (left) and water (right) intake of the mice in the 4 days before sacrifice (food: F(1,70) = 0.5572, P = 0.458; water: F(1,50) = 0.1192, P = 0.73; two-way RM ANOVA with Sidak’s post hoc test). **f**. Blood glucose levels at different time points (F(1,38) = 29.7, P = 3.24 × 10^−6^, *P = 0.013, **P = 0.0056, two-way RM ANOVA with Sidak’s post hoc test). **g**. Inguinal fat mass (t = 2.78, *P = 0.0195, t test). **h**. Quadriceps muscle mass (t = 4.65, ***P = 0.0009, t test). **i**. Tumor mass (t = 7, ****P = 0.000037, t test). **j**. Spleen mass (t = 3.43, **P = 0.0064, t test).

